## Supplementary information for "Diverse stem-chondrichthyan oral structures and evidence for an independently acquired acanthodid dentition"

**Supplementary table 1:** details on specimens scanned

| Taxon | Specimen number | Scan Location | Scanner Model | Voxel size (μm) | Current (μA) | Power (kV) | Filter | Projections |
|---|---|---|---|---|---|---|---|---|
| *Acanthodes* | NHMUK PV P.8065 | Department of Life Sciences, Bristol University | Nikon XT H 225 ST | 44.9 | 165 | 215 | 0.1 mm Sn | 3142 |
| *Acanthodopsis* | NHMUK PV P.10383 | Department of Life Sciences, Bristol University | Nikon XT H 225 ST | 22.6 | 92 | 180 | None | 3142 |
| *Atopacanthus* | NHMUK PV P.10978 | Imaging and analysis centre, NHMUK | Metris X-Tek HMX ST 225 | 19.508 | 154 | 130 | 0.1 mm Cu | 3142 |
| *Ischnacanthus* | NHMUK PV P.40124 | Department of Life Sciences, Bristol University | Nikon XT H 225 ST | 24.6 | 105 | 222 | 0.5 mm Cu | 3142 |
| *Taemasacanthus* | NHMUK PV P.33706 | Imaging and analysis centre, NHMUK | Metris X-Tek HMX ST 225 | 17.3 | 131 | 130 | 0.1 mm Cu | 3142 |

### Notes on phylogenetic analysis

Changes made to the phylogenetic matrix of Dearden *et al* are detailed below.

#### Taxon addition: *Acanthodopsis*

Of the taxa in our list *Acanthodopsis* is the only one with sufficient characters to code into the wider matrix. Scorings were taken from our data and Burrow (2004).

#### Character addition (character 268): Mandibular splint:

(0) absent, (1) present.

As discussed in the main text a mandibular splint, defined as a smooth dermal bone along the ventrolateral margin of the jaw, is present in acanthodid acanthodians.

#### Character addition (character 269): Dental plates with smooth occlusal surface:

(0) absent, (1) present

This is taken from character 261 of Burrow *et al.* (2016). The dermal jaw plates of putative diplacanthid stem-chondrichthyans are smoothly convex bones with a completely unornamented surface, as discussed in the main text. This character is inapplicable for taxa without dermal jaw plates.

#### Character addition (character 270): Expanded mandibular symphysis at anterior tip of Meckel's cartilage:

(0) absent, (1) present

As noted in the main text some acanthodids have an expanded region at the anterior end of the Meckelian cartilage. *Acanthodopsis* exhibits the same morphology as *Acanthodes* (RPD, pers. obs.). *Cheiracanthus* and *Homalacanthus* also appear to share the same form (see Burrow et al., 2020 fig 7.1; Miles, 1966 fig 16B). In *Halimacanthodes* the symphysis is rounded, but otherwise has the same morphology (Burrow et al., 2012).

*Ischnacanthus* and close relatives appear to lack this morphology (Blais et al., 2015; Burrow et al., 2018), as does *Promesacanthus* (Hanke and Wilson, 2004). Meckel's cartilages in most other acanthodian-grade stem-chondrichthyans are poorly known, but it appears to be absent in stem-chondrichthyans *Gladbachus*, *Doliodus*, and *Pucapampella* (Coates et al., 2018; Maisey et al., 2019; Miller et al., 2003).

This character is scored inapplicable for taxa with a fused mandibular symphysis.

#### Character addition (character 271): Scapular Process has expanded waist:

(0) absent, (1) present

In *Acanthodes* and *Acanthodopsis* the scapular process bulges outwards about halfway up its length (Burrow, 2004; Miles, 1973). This distinctive morphology is absent in other chondrichthyans and in other 'acanthodians', and notably absent in *Halimacanthodes*, which is otherwise anatomically similar to *Acanthodes* (Burrow et al., 2012). This character is contingent on having a scapular process.

#### Coding change: (character 114) Dorsal process on lower dermal jaw plate

This character was originally erected to capture a similar dorsal process between the "Meckelian elements" of *Tetanopsyrus* and *Uraniacanthus* (Brazeau, 2009; Hanke and

Wilson, 2004). This character has been redefined to explicitly refer to a lower dermal jaw plate rather than a "Meckelian bone or cartilage". It is inapplicable for taxa without a dermal jaw plate of the lower jaw.

Dorsal processes of the dermal jaw plate are only present in *Uraniacanthus* (Hanke and Davis, 2008; Newman et al., 2012) and *Culmacanthus* (Burrow and Young, 2012). *Tetanopsyrus* is coded 0 as its "dorsal process" is of the Meckelian element - the lower dental plate has no process (Hanke et al., 2001). *Diplacanthus* has in the past been scored present for this character (Davis et al., 2012): this is likely due to observation of the articular part of the Meckelian cartilage in NMS 1891.92.334 (Watson, 1937): its occlusal plate lacks a dorsal process (Burrow et al., 2016) and so it is scored 0. This character is sometimes coded present in the problematic gnathostome *Ramirosuarezia* (e.g. Coates *et al.* (2018)) on the basis of a dorsal process on what is interpreted as its Meckel's cartilage (Pradel et al., 2009). However this is non-homologous with the dermal plate of diplacanthids, evidenced by the fact that it bears tooth plates, and we score *Ramirosuarezia* is here scored 0 for this character.

### Coding change: Jaw dermal plates (Character 94)

The scoring for *Culmacanthus* was changed from ? to 1 on the basis of it's having occlusal plates (Burrow and Young, 2012)

#### Coding change: Gnathal plates mesial (Character 95)

The scoring for *Culmacanthus* was changed from 0 to ? on the basis of its occlusal plates not being preserved in articulation with jaws (Burrow and Young, 2012).

#### Coding change: Tooth whorls (Character 84)

The scoring for *Lupopsyrus* and *Halimacanthodes* was changed from 0 to – as in other toothless taxa so as to be consistent with this character's contingency on having teeth.

#### Coding change: Oral tubercles on jaws (Character 82)

*Tetanopsyrus* is described as having teeth by Hanke, Davis and Wilson (2001) on the basis of an isolated pair of jaws (Hanke et al., 2001 fig. 3B) found close to the articulated *Tetanopsyrus lindoei* specimen UALVP 32571. These jaws are preserved in lingual view and comprise high-profiled Meckel's cartilages with a what are interpreted as smooth occlusal plates with a row of tooth-like tubercles on the lingual face. These isolated jaws do not belong to UALVP 32571 itself, which has lower jaws preserved in articulation (Hanke et al., 2001 fig. 3A) . They are attributed to *Tetanopsyrus* presumably on the basis of having a "smooth" occlusal plate (which is not seen in lingual view in any other published specimen) as well as a similarly profiled Meckelian cartilage. However, no clear characters place the jaws in the genus Tetanopsyrus.

While these jaws may prove to belong to *Tetanopsyrus*, here we have taken a conservative approach, and have code *Tetanopsyrus* ? for the presence of teeth (character 82) and for all contingent characters.

**Supplementary figure 1.** Vasculature in dentigerous jaw bones in *Taemasacanthus* NHM PV P.33706 (a) medial view, (b) close-up of boxed area in (a), (c) lateral view, and (d) ventral view. *Atopacanthus* NHMUK PV P.10978 in (e) medial view, (f) close up of boxed area in (e) and (g) lateral view. *Ischnacanthus* NHMUK PV P.40124 in (h) lateral, and (i) medial view. Abbreviations: lat.t.r., lateral tooth row; ling.t.r, lingual tooth row; ling. vasc, lingual vasculature; med.t.r., medial tooth row; mes.ri, mesial ridge; pol. vasc, polarised vasculature; pulp. cav., pulp cavity; rad. vasc., radial vasculature; young.t, out-of-order youngest tooth. Scale bars = 5 mm.

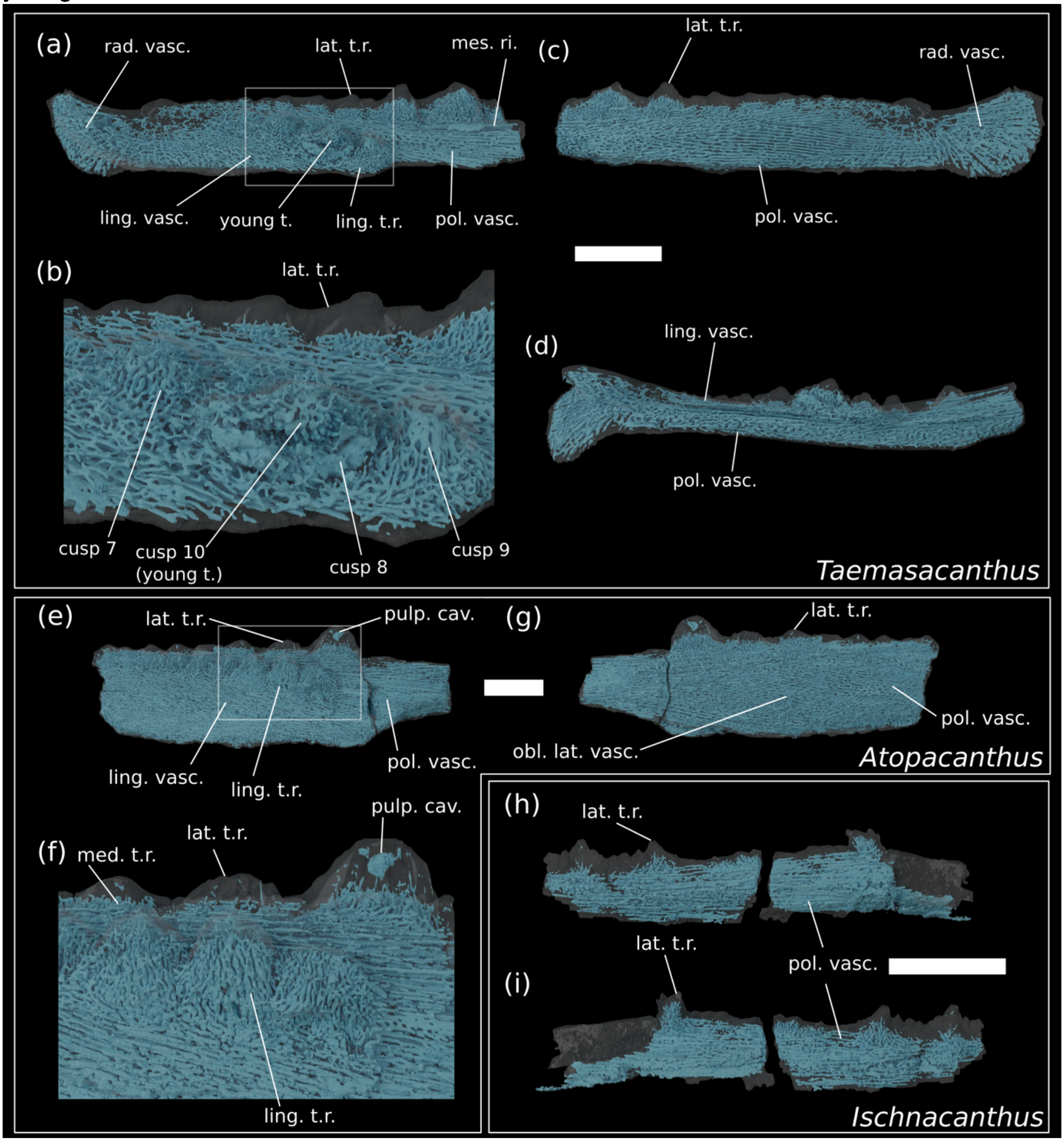

**Supplementary figure 2.** Complete results of the parsimony analysis in main text Fig. 5. (a) strict consensus tree with Bremer support values on dichotomous nodes. (b) Adams consensus.

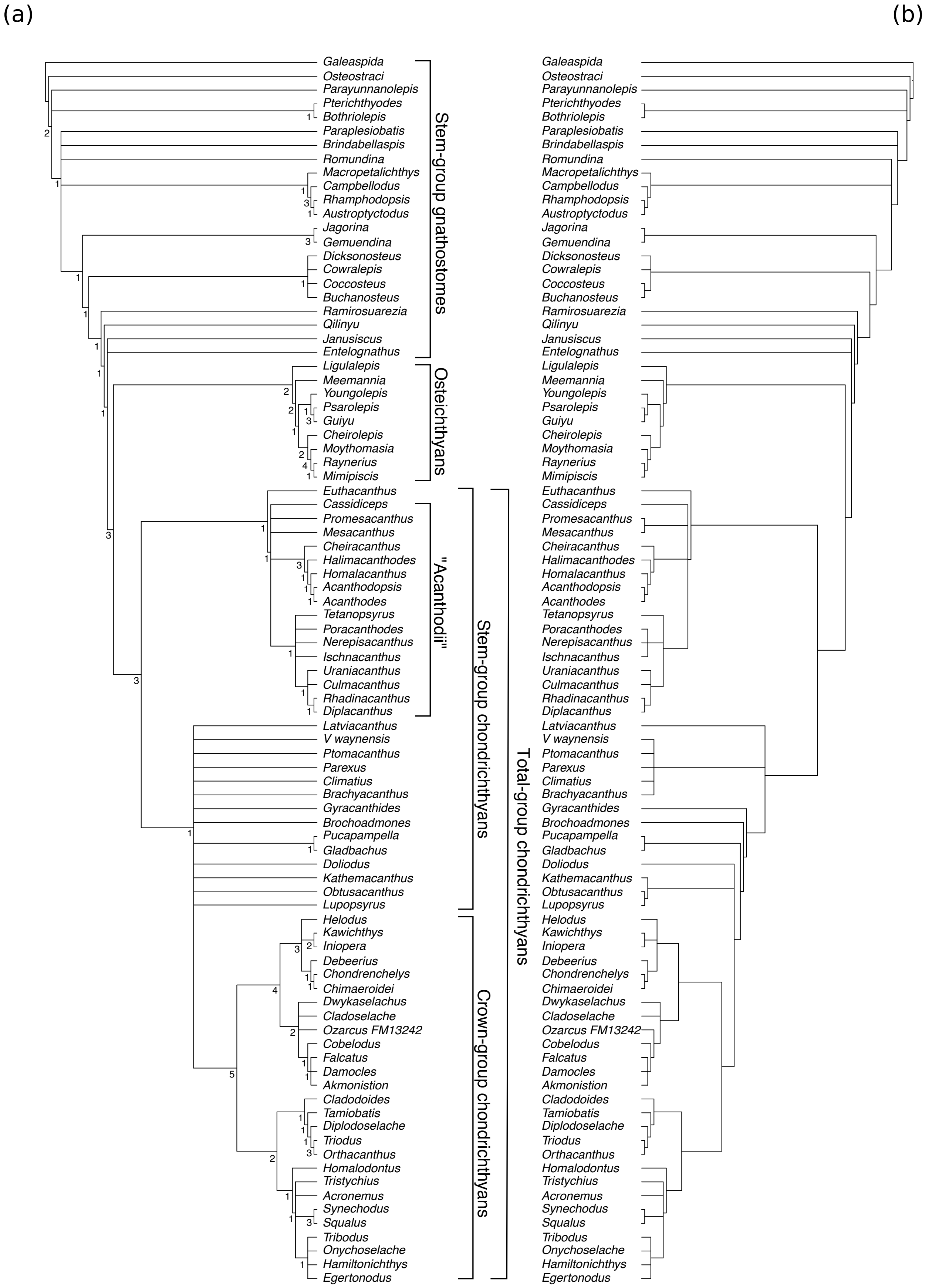

**Supplementary figure 3.** Majority rule consensus tree for the Bayesian analysis of the phylogenetic dataset. Nodes are labelled with posterior probabilities.

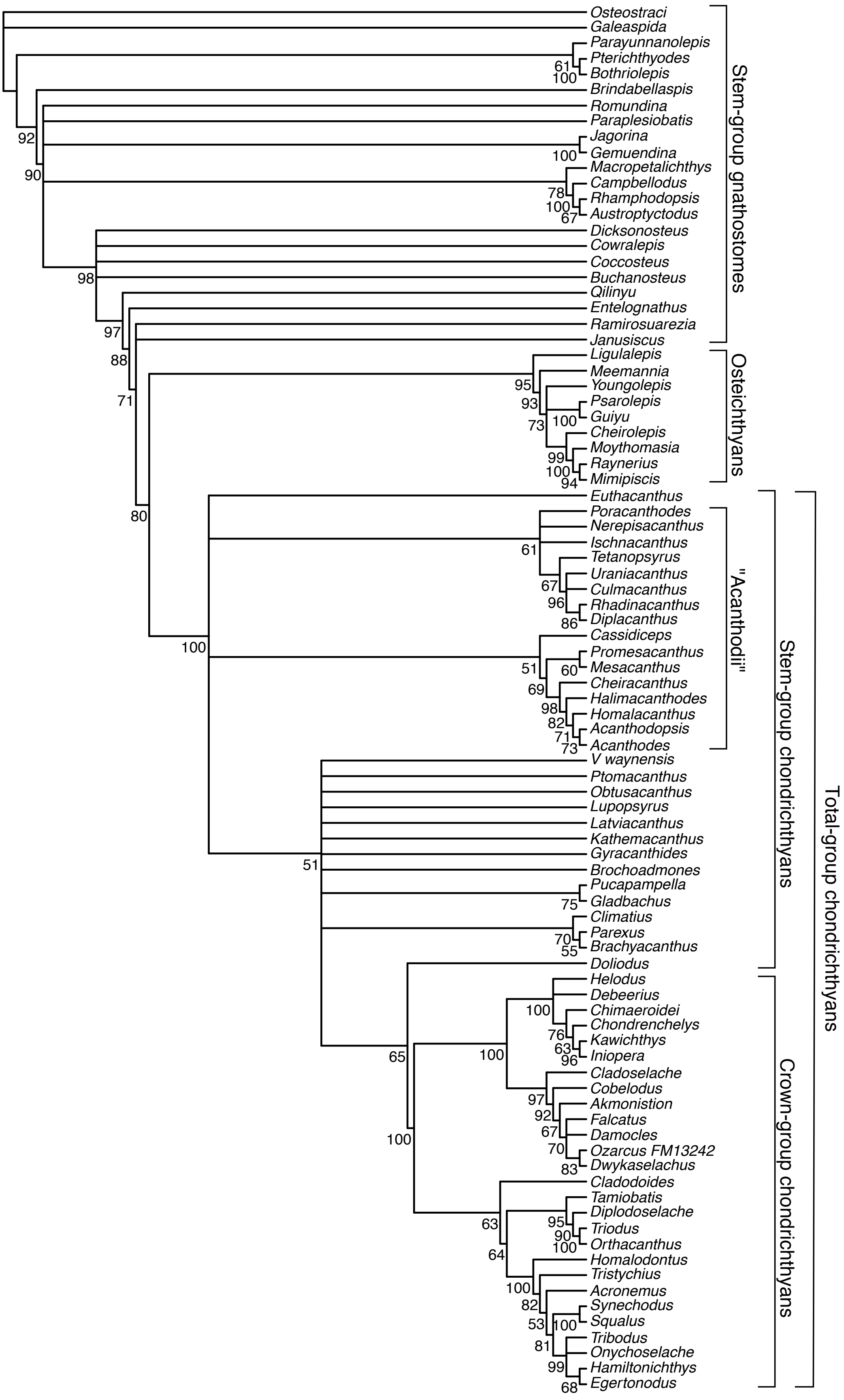
